# Discovery and Characterization of a Hydroxypyridone-3-carboxamide Analog as an Antiviral Lead against Orthopoxviruses

**DOI:** 10.64898/2025.12.18.695249

**Authors:** Anil Pant, Ajit Jagtap, Jiashu Xie, Djamal Brahim Belhaouari, Wei Xie, Jacob P. Mahoney, Lara Dsouza, Ziyue Wang, Zeinab R. Aboezz, Tibor Farkas, Zhengqiang Wang, Zhilong Yang

**Affiliations:** Department of Veterinary Pathobiology, College of Veterinary Medicine & Biomedical Sciences, Texas A&M University, College Station, TX 77843; Center for Drug Design, College of Pharmacy, University of Minnesota, Minneapolis, MN 55455

**Keywords:** Orthopoxvirus, Mpox, Antiviral, Organoids, Enteroids, *ex vivo* lung tissue, Hydroxypyridone-3-carboxamide

## Abstract

Orthopoxviruses remain a persistent global health concern due to the ongoing circulation of mpox, the possibility of the re-emergence of smallpox, and the threats posed by many poxviruses that infect animals and/or humans. The limited availability of antiviral drugs, the unproven efficacy in humans, and the emergence of resistant mutants underscore the need for new and better therapeutics. In this study, we identify and characterize ZW-2038, a hydroxypyridone-3-carboxamide analog, as an antiviral compound against vaccinia virus (VACV), monkeypox virus (MPXV), and cowpox virus (CPXV). Discovered through a focused in-house small-molecule screen, ZW-2038 exhibited low micromolar potency and high selectivity in primary human fibroblasts. The compound also reduced viral replication under physiomimetic conditions including human and monkey intestinal organoids (enteroids) and *ex vivo* mouse lung tissue models. Mechanistically, ZW-2038 suppresses VACV DNA replication and downstream post-replicative gene expression, albeit without inhibiting MPXV resolvase (Mpr) *in vitro*. These findings, along with *in vitro* safety profiling and mice pharmacokinetics studies, characterize ZW-2038 as a promising yet suboptimal antiviral lead against orthopoxviruses warranting future development.

## Introduction

Poxviruses continue to pose real threats to human and animal health. Although declared eradicated in 1980, smallpox, one of the most devastating diseases in human history, remains a concern due to the possibility of accidental release, deliberate misuse, or *de novo* biosynthesis (1, 2). The end of routine vaccination and waning immunity across the global population creates the conditions for rapid spread if smallpox were ever to re-emerge (3). The Mpox outbreaks caused by monkeypox virus (MPXV) in recent years has shown how vulnerable we remain to poxviruses. The World Health Organization (WHO) has issued two declarations of Public Health Emergency of International Concern (PHEIC) related to mpox in the past three years: the first in 2022 in response to the widespread multinational outbreak caused by Clade IIb, and a second in August 2024 following the rapid expansion of the more virulent Clade I b outbreak in the Democratic Republic of the Congo and adjacent regions (4, 5). The rapid global spread of MPXV during the 2022 outbreaks, with reported cases in more than 142 countries and territories, including over 115 countries where it was not previously reported, has raised serious concerns about the potential for a poxvirus pandemic (6, 7). MPXV infections can be severe, especially in immunocompromised individuals, and the virus has shown ability to adapt to new routes of transmission and spread systemically (8–11). Animal poxviruses such as lumpy skin disease virus, capripoxviruses, and swinepox lead to disease outbreaks in cattle, sheep, goats, and pigs, resulting in billions of dollars in economic losses each year (12, 13). These infections reduce productivity, disrupt trade, and threaten the livelihoods of farmers. Moreover, they create opportunities for cross-species transmission, increasing the risk of novel poxviruses entering human populations.

There is a clear and urgent need to develop new antivirals against poxviruses, as the current therapeutic options are limited and not consistently effective. Tecovirimat and brincidofovir are the two antiviral drugs that have been approved by the FDA for strategic stockpiling against smallpox. Although tecovirimat is a promising and most commonly used option, it was developed and approved under the Animal Rule (14, 15). More recently, two randomized, placebo-controlled trials have shown that tecovirimat does not improve clinical outcomes in children or adults infected with clade I MPXV, or in adults infected with clade II MPXV (16–19). Cidofovir, approved for treating human cytomegalovirus (HCMV), and its prodrug brincidofovir, have occasionally been used because they target viral DNA polymerase, but their efficacy against mpox in humans remains unknown (20), and clinical use of cidofovir is associated with severe side effects (21–23). As orthopoxviruses, including mpox and related zoonotic poxviruses, continue to expand into new geographic regions and host populations, there is a critical need to develop chemically novel and mechanistically distinct antivirals that are accessible to global populations.

Toward this end, we phenotypically screened an in-house small molecule library consisting of representative compounds from various chemotypes previously synthesized to target other viruses. Using the *Gaussia* luciferase-based reporter assay that we have established in our lab, this study identified compound ZW-2038, a hydroxypyridone-3-carboxamide (HPCAm) analog as a strong hit against VACV. ZW-2038 was previously reported as a potent antiviral compound against HCMV (24). In the current studies, we found that ZW-2038 effectively and selectively suppresses the replication of vaccinia virus (VACV), cowpox virus (CPXV), and MPXV in primary human cells, human and non-human primate enteroids and *ex vivo* mouse lung tissues. Mechanistically, we found that ZW-2038 suppresses VACV replication by inhibiting VACV DNA replication and subsequent post-replicative gene expression. However, in an *in vitro* assay ZW-2038 did not inhibit MPXV resolvase (Mpr), an enzyme involved in poxvirus genome replication (25), despite bearing a metal chelating pharmacophore shared among Mpr inhibitors. Pharmacokinetics (PK) studies in mice produced low oral bioavailability, presumably due to poor aqueous solubility as shown in previous ADME studies (24). In the drug safety hERG assay, ZW-2038 produced marginal inhibition, suggesting low hERG liability. Collectively, these studies characterize ZW-2038 as a promising antiviral candidate with clear potency and ADME goals for further optimization.

## Materials and Methods

### Cells and viruses

Primary human foreskin fibroblasts (HFFs, kind gift from Dr. Nicholas Wallace, Kansas State University) and BS-C-1 Cells (ATCC CCL-26) were maintained in DMEM or EMEM supplemented with 10% fetal bovine serum (FBS; VWR), 2 mM L-glutamine (VWR), 100 U/mL penicillin, and 100 μg/mL streptomycin (VWR) at 37 °C in a humidified incubator with 5 % CO2.

Vaccinia virus (VACV), Cowpox virus (CPXV, Brighton Red strain), and a 2022 isolate of mpox virus (MPXV-MA001, GenBank: ON563414.3) were used in this study. We utilized recombinant VACVs that express *Gaussia* luciferase under the control of early (C11R, vEGluc), intermediate (G8R, vIGluc), or late (F17R, vLGluc) viral promoters, as previously described (26). These reporters allow for rapid and real-time detection of gene expression since *Gaussia* luciferase is secreted into the culture medium upon synthesis, eliminating the need for cell lysis (27). All virus propagation, purification, and infection procedures were performed using standard published protocols (28).

### *Ex vivo* mouse lung tissue infection assays

The collection, infection, and titration of VACV in *ex vivo* mouse lung tissues was performed as described previously (29). Briefly, lung tissue was collected from C57BL/6 (B6) mice immediately following euthanasia under humane conditions. After rinsing with phosphate-buffered saline, lungs were sliced into thin sections using a McIlwain tissue chopper. Equal weights of lung slices (typically 4–5 per sample) were placed into wells of a 24-well tissue culture plate. The slices were then infected with 1 × 10⁵ PFU of vLGluc VACV, and treated with either 10 μM ZW-2038, 40 μg/mL AraC, or vehicle control. One-hour post-infection, the inoculum was removed and replaced with fresh media containing the corresponding treatment. After indicated time points, the tissues were homogenized, and the samples were freeze-thawed for three times. The virus titers in the samples were measured using a plaque assay as described previously. All experiments were conducted using three independent biological replicates. The mouse work was approved by the Institutional Biosafety Committee at Texas A&M University (Approval of Animal Use Protocol IACUC, 2022-0224).

### Enteroid culture and infection

Human jejunum organoids (J2) and Rhesus macaque colon (NC67) organoids (enteroids) were cultured in 3D Matrigel droplets and maintained as described in previous studies (30, 31). To initiate cultures, frozen stock of enteroids was thawed and resuspended in 10 mL of ice-cold complete medium without growth factors (CMGF-), consisting of Advanced DMEM/F-12 supplemented with 1% HEPES (1 M), 1% GlutaMAX, and 1% penicillin-streptomycin (10,000 U/mL). Cells were gently pelleted by centrifugation (300 × g, 5 min, 4 °C), mixed with Corning GFR Matrigel, and plated as 20 μL domes in 24-well tissue culture plates. Plates were kept at room temperature for 1 minute to allow Matrigel to set, then inverted and incubated at 37 °C for 10 minutes before adding warm complete medium with growth factors (CMGF+). All enteroids were passaged every 7–10 days and used for infection studies when cultures reached appropriate density and morphology.

Complete growth media (CMGF+) consisted of L-WRN conditioned media and an equal volume of CMGF-media, and supplemented with 10% FBS, recombinant mouse EGF (50 ng/mL, Invitrogen), nicotinamide (10 mM, Sigma), gastrin I (10 nM, Sigma), A-83-01 (500 nM, Tocris), SB202190 (10 μM, Sigma), B27 and N2 supplements (1× each, Invitrogen), and N-acetylcysteine (1 mM, Sigma). The L-WRN conditioned media containing Wnt-3A, R-spondin, and Noggin was prepared in house using L-WRN cells (ATCC CRL-3276). For infections, enteroids were harvested, washed, and mechanically dissociated into small fragments to allow viral access. vLGluc was added at indicated multiplicity of infection, and infections were carried out in suspension for 1 hour at 37 °C with gentle rocking. Following infection, enteroids were washed and re-embedded in Matrigel. Fresh media with or without test compounds was added, and supernatants were collected at defined time points to measure luciferase activity as a readout of viral replication.

### Titration of viruses

VACV, CPXV, and MPXV titers were calculated by plaque assay using the protocol described elsewhere (28). Briefly, BS-C-1 cells were cultured in 12-well plates were overlayed with 10-fold diluted virus samples collected at indicated time post treatment with vehicle or indicated compounds and incubated in EMEM media with 2.5% FBS. At 1 hour post infection (hpi) the media was changed to EMEM 10% FBS and 0.5% methyl cellulose and incubated for 48 h. The cells were then stained with 0.1% crystal violet for 15 min and washed with water before counting the number of plaques.

### Chemicals

Cytarabine (AraC) was purchased from Sigma-Aldrich. Tecovirimat was purchased from TargetMol and was dissolved in DMSO for all experiments. ZW-2038 was synthesized according to reported method (24) and fully characterized with 1H, 13C and 19F NMR, HRMS and HPLC (See the Supplementary information for the synthetic scheme and analytical data). The purity is 97% as determined on analytical HPLC under these conditions: flow rate, 1.0 mL/min; solvent A, 0.1% formic acid in water; solvent B, 0.1% formic acid in acetonitrile; gradient (B, %): 0–6 min (5–100), 6–8 min (100), 8–9 min (100–5). The chemicals needed for enteroid culture are listed in the respective section.

### *Gaussia* luciferase assay

Cells, tissues, or enteroids were infected with recombinant VACV encoding secreted *Gaussia* luciferase under early, intermediate, or late promoters at indicated multiplicity of infection. The activities of *Gaussia* luciferase in culture medium were measured at indicated hpi using a Pierce *Gaussia* luciferase flash assay kit (Thermo Scientific) and a GloMax Luminometer (Promega).

### Quantitative real-time PCR

Total DNA was extracted from the cells using the EZNA Blood DNA Kit. Relative viral DNA levels were quantified by CFX96 real-time PCR instrument (Bio-Rad, Hercules, CA) using All-in-oneTM 2 × qPCR mix (GeneCopoeia) using primers against the VACV C11 gene and normalized to 18S rRNA levels as an internal control. The sequences for the primers used are as listed below:

C11p FW: AAACACACACTGAGAAACAGCATAAA.

C11p Rev: ACTATCGGCGAATGATCTGATTATC.

18S rRNA FW: CGA TGC TCT TAG CTG AGT GT.

18S rRNA Rev: GGT CCA AGA ATT TCA CCT CT.

### Cell viability, CC_50_, and EC_50_ determination

The viability of the cells was measured using the trypan blue exclusion method as described previously (32). Briefly, cells were treated with DMSO or the indicated compounds at indicated concentrations and counted using a TC20 automated cell counter (Bio-Rad) after staining with equal volume of 4% trypan blue.

The 50% cytotoxic concentration (CC_50_) of the compounds was determined using the CCK-8 assay according to the manufacturer’s instructions. Briefly, HFFs were grown for 24 h in 96-well plates. Then the cells were treated with DMSO or serially diluted concentrations of indicated compounds. After 24 h incubation, 10 µL of CCK-8 reagent was added to each well, and the cells were incubated for 3 h in a 37 °C incubator. For each well, the absorbance at 450 nm was measured using a BioTek Cytation 5 imaging reader (Agilent Technologies). CC₅₀ values were calculated by fitting the data to a non-linear regression model (“log(inhibitor) vs. normalized response-variable slope”) in GraphPad Prism (version 10.2.3).

To determine the half-maximal effective concentration (EC₅₀) of the compounds for inhibiting viral replication, we used a recombinant VACV that expresses secreted *Gaussia* luciferase from a viral late promoter (vLGluc), following established protocols (29, 33, 34). HFFs were seeded in 96-well plates and infected with vLGluc at MOI of 0.01 in the presence of either DMSO or serial dilutions of the test compounds. At 24 h post-infection, *Gaussia* luciferase activity in the culture supernatant was measured using a luminometer. EC₅₀ values were calculated by fitting the data to a non-linear regression model (“log(inhibitor) vs. normalized response-variable slope”) in GraphPad Prism (version 10.2.3).

### Mpr assays

Mpr with an N-terminal maltose binding protein fusion and C-terminal 6xHis tag was expressed in *Escherichia coli* and purified as previously described (25). Mpr activity was monitored using a FRET assay in which cleavage of a fluorophore- and quencher-labeled DNA substrate by Mpr produces a product with increased fluorescence intensity (25). Reactions were performed in black 384-well plates (Corning) using a buffer containing 20 mM Tris (pH 8.5 at 22 °C), 50 mM NaCl, 10 mM MgCl_2_, 5% glycerol, 5 mM β-ME, 0.0025% v/v Tween 20, 0.25% v/v DMSO, along with the indicated compounds at 50 µM final concentration. Mpr (10 nM) was incubated with compound or DMSO for 15 min at 37 °C. Substrate (5 nM) was added and fluorescence intensity (excitation: 485 nm; emission: 535 nm) was monitored for 15 min at 37 °C using a SpectraMax i3 reader (Molecular Devices). The rate of fluorescence increase was determined from the linear portion of the data using GraphPad Prism 10.

### hERG assay

The Predictor hERG Fluorescence Polarization Assay (PV5365, ThermoFisher Scientific) was used to assess the hERG channel binding activity of the test compounds. Stock solutions were prepared at 100-fold in DMSO (Sigma Aldrich) and subsequently diluted to 4-fold in assay buffer. Appropriate volumes of assay buffer, 4-fold test compound solutions, or positive control (E-4031, supplied with kit) were dispensed into designated wells of a 384-well plate (Corning).

Following this, 10 μL of 2-fold Predictor hERG Membranes were added to each well, followed by 5 μL of 4-fold Predictor hERG Tracer Red. Plates were incubated for 3 hours at room temperature, after which fluorescence polarization was measured using a Tecan Spark™ 10M microplate reader (excitation: 534 nm; emission: 574 nm).

Concentration response curves were generated and fitted using a four-parameter sigmoidal dose–response model in GraphPad Prism version 10.6.1 (GraphPad Software, Boston, MA)

### Mice PK

Oral bioavailability of ZW-2038 was assessed in 7-week-old male CD-1 mice (strain code 022), obtained from Charles River Laboratories (Wilmington, MA, USA). Animals were housed and cared for at the AAALAC-accredited Research Animal Resources facility of the University of Minnesota. All procedures were approved by the University’s Institutional Animal Care and Use Committee (IACUC; protocol ID 2110-39488A) and conducted in accordance with IACUC policies and institutional animal welfare guidelines. Briefly, the compound was administered via two routes: tail vein intravenous (IV, 2 mg/kg) and oral gavage (PO, 20 mg/kg), with four mice randomly assigned to each route. Blood samples (∼20 µL per time point) were serially collected via saphenous vein puncture into EDTA-fortified tubes at 5, 15, and 30 min, and at 1, 2, 4, 8 (IV: 7), and 24 (IV: 23) h post-dose, then centrifuged at 3000 rpm for 15 min to obtain plasma. Plasma concentrations were quantified using LC/MS/MS with a matrix-matched calibration curve. Concentration-time data were analyzed using Phoenix WinNonlin (version 8.5.2.4; Certara USA, Inc., Radnor, PA, USA) employing non-compartmental modeling. Oral bioavailability (F%) was calculated by comparing the area under the curve (AUC) for oral and intravenous administration according to Eq. (1):

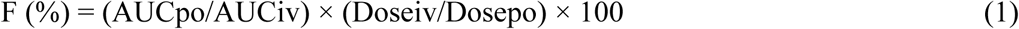

### Statistical Analyses

All data are shown as the average with error bars representing standard deviation (SD), based on at least three independent experiments unless otherwise noted. We used GraphPad Prism (version 10.2.3) and Microsoft Excel (version 16.43) for the analyses. To compare two groups, we used a two-tailed paired t-test. For comparing more than two groups, we used one-way ANOVA followed by either Tukey’s or Dunnett’s post hoc tests. When comparing across two different factors, we used two-way ANOVA followed by Tukey’s test. Statistical significance was marked as: Not significant (ns): P > 0.05; *P ≤ 0.05; **P ≤ 0.01; ***P ≤ 0.001; ****P ≤ 0.0001.

## Results

### Focused library screening identifies ZW-2038 as a potent antiviral candidate against VACV

We screened a focused in-house library of 398 in-house small molecules for their ability to inhibit the replication of VACV (**Fig. 1A**). The compounds were selected from a larger collection to represent many chemotypes previously synthesized to target other viruses. Primary Human Foreskin Fibroblasts (HFFs) were seeded in 96-well plates and after overnight culture, they were infected with vLGluc at MOI of 0.01. After 1 hpi, the media was replaced with vehicle or different compounds at 10μM concentration. The supernatant was transferred to clean, flat-bottom, opaque assay plates after 24 hpi and the *Gaussia* luciferase activity was measured.

**Fig 1.**
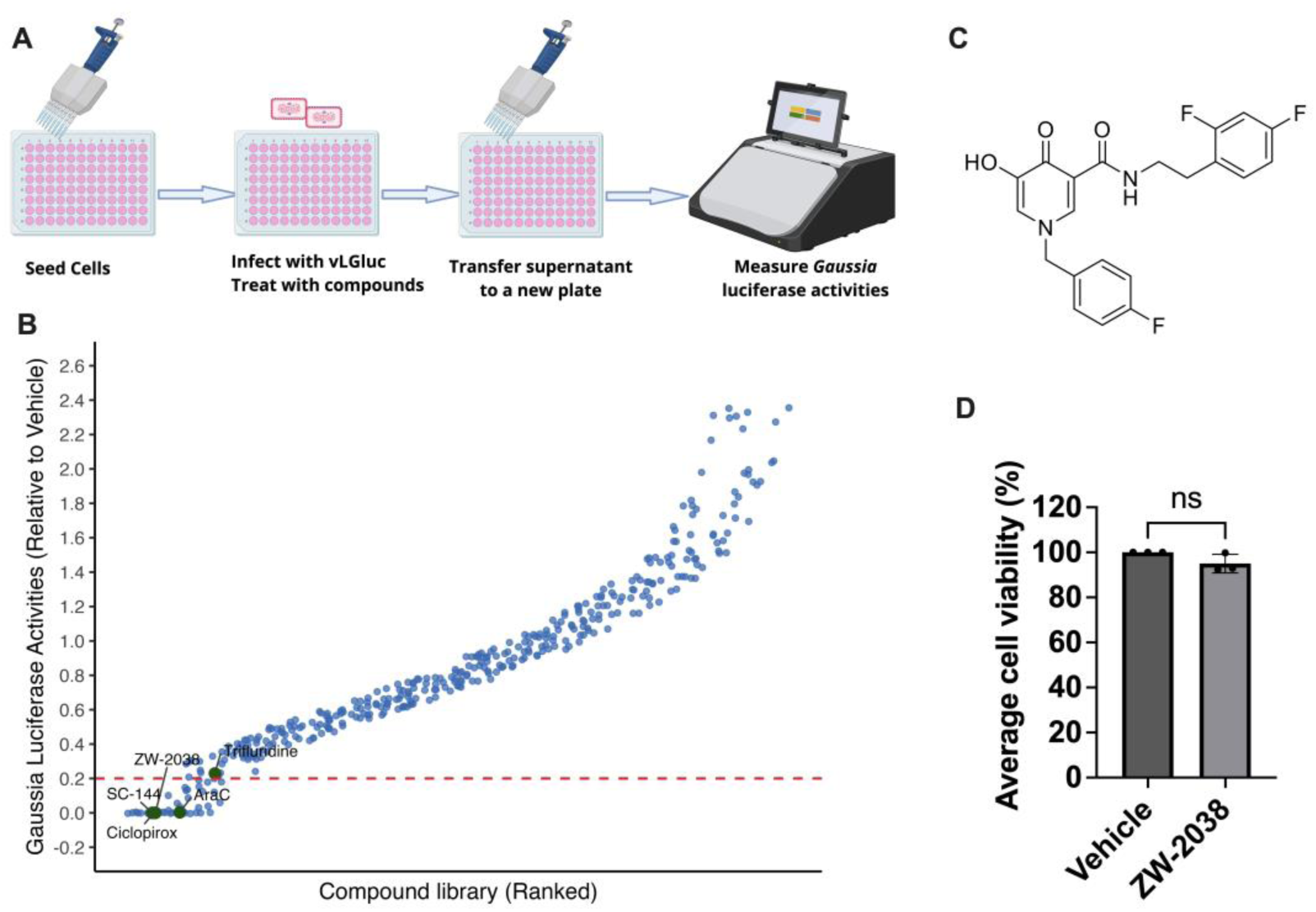
Focused library screening identifies ZW-2038 as a potent antiviral candidate. **(A)** Schematic of the vLGLuc reporter assay workflow. Cells were seeded in 96-well plates, infected with recombinant VACV expressing *Gaussia* luciferase (vLGLuc), and treated with vehicle or test compounds at 10 µM concentration. After incubation, cell culture supernatants were transferred to a new plate, and luciferase activity was measured to quantify viral replication. **(B)** Small molecule compound library screening identifies ZW-2038 among the top hits that inhibit VACV replication. A library of 398 small-molecule compounds was screened for anti-VACV activity using a *Gaussia* luciferase reporter assay. Luciferase activity in compound-treated supernatants was measured and normalized to vehicle-treated controls. Compounds were ranked according to their relative antiviral activity, with the red dotted line indicating an empirical threshold of 80% or greater reduction in viral replication compared to vehicle control. ZW-2038, ciclopirox, and several other known and novel antivirals clustered among the top candidates. **(C)** Chemical structure of ZW-2038. **(D)** Effect of ZW-2038 treatment on host cell viability. HFFs were treated with vehicle or ZW-2038 (10 µM) for 48 h, and cell viability was measured using the trypan blue assay. Data are presented as mean ± SD (n = 3) and analyzed by unpaired t-test; ns = not significant.

ZW-2038, an HPCAm analog (**Fig. 1C**), emerged as a top hit that showed strong inhibition of *Gaussia* luciferase activity in the infected samples with more than 99% reduction (**Fig. 1B**). The reduction was comparable to that of Ciclopirox and SC144, which have been characterized as efficient inhibitors of VACV replication in our previous work (29, 34). Trypan blue staining revealed no significant difference in the cell viability in the ZW-2038 treated cells compared to vehicle treatment (**Fig. 1D**). Overall, the screening identified ZW-2038 as a promising hit suppressing VACV replication without affecting HFF viability.

### ZW-2038 is a potent and selective lead of VACV replication inhibitor in primary HFFs

To validate the antiviral activity of ZW-2038, we infected HFFs with VACV at both high (MOI = 2) and low (MOI = 0.01) MOIs in the presence of 10 μM ZW-2038. Cytarabine (AraC) was used as a positive control. Using plaque assays, which is considered the gold standard for quantifying infectious virus (35), we found that ZW-2038 reduced VACV titers by over 295-fold at MOI 2 (**Fig. 2A**) and 2300-fold at MOI 0.01 (**Fig. 2B**) compared to the DMSO-treated controls. The extent of suppression was similar to that observed with AraC at both MOIs.

**Fig 2.**
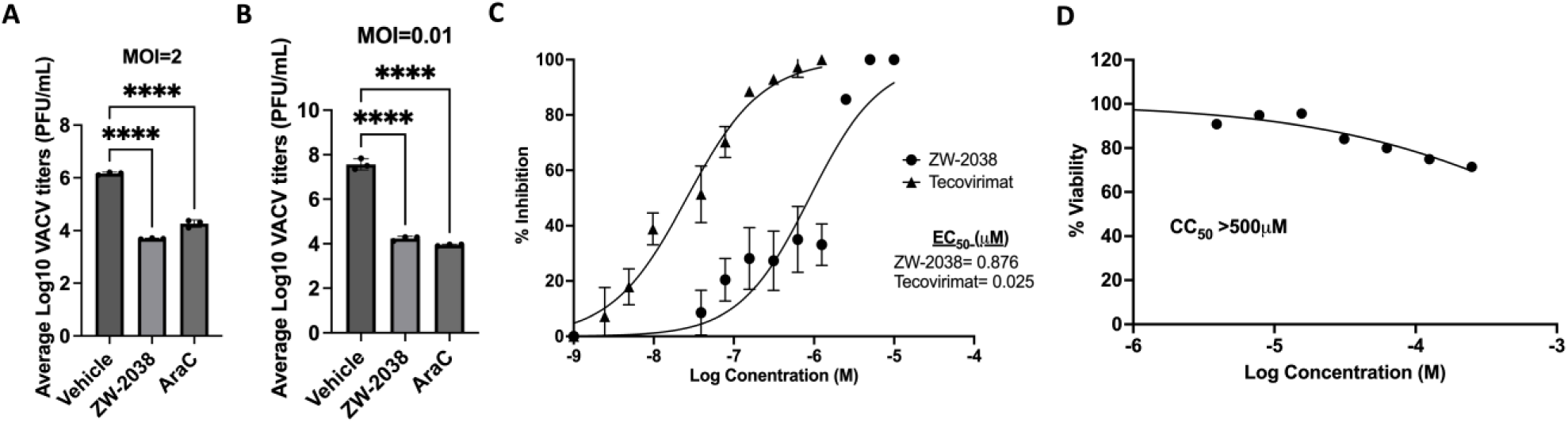
ZW-2038 effectively and selectively suppresses VACV replication at low micromolar concentration. **(A-B)** Primary human foreskin fibroblasts were infected with VACV at high (MOI = 2, **A)** or low (MOI = 0.01, **B)** multiplicity of infection and treated with 10 μM ZW-2038, 40 μg/mL AraC (positive control), or vehicle control. Cell pellets were collected at 24 hpi (A) and 48 hpi (B) and infectious virus titers were quantified by plaque assay. **(C)** HFFs were infected with MOI = 0.01 of vLGluc in the presence of vehicle or a series of concentration of ZW-2038 or tecovirimat. *Gaussia luciferase* activities were measured at 24 hpi to determine the EC_50_ value. Non-linear regression was used to calculate EC_50_ values. **(D)** HFFs were treated with vehicle or a series of concentration of ZW-2038. CCK8 assay was performed at 24 h post treatment to calculate the CC_50_ value. All data represent mean ± SD of at least three independent biological replicates. Statistical analysis was performed using one-way ANOVA with Dunnett’s post hoc test. ****p<0.0001.

Next, we examined the antiviral potency of ZW-2038 by measuring its half-maximal effective concentration (EC_50_) using a luciferase-based assay with recombinant VACV expressing *Gaussia* luciferase under the viral late promoter (vLGluc). We have extensively optimized this luciferase system for quantifying viral replication in live cell settings and established it as a valid method for EC_50_ measurements (27, 29, 33, 34). At MOI of 0.01 and 24 h infection incubation time, the EC_50_ of ZW-2038 was calculated to be 0.876 µM in HFFs (**Fig. 2C**). Using the same MOI and incubation time, we determined the EC_50_ of tecovirimat was 0.025 µM, which is comparable to what was reported in our previous study (34). ZW-2038 exhibited minimal cytotoxicity in HFFs with CC_50_ of over 500 µM (and selective index of over 1480) at 24 hours post treatment (**Fig. 2D**) indicating this compound does not cause cytotoxicity in HFFs even at high concentrations. Overall, these results show that ZW-2038 is a potent and selective inhibitor of VACV replication in cultured cells.

### ZW-2038 suppresses VACV DNA replication and downstream gene expression but not early gene replication

VACV replication progresses through sequential stages, beginning with entry and early gene transcription, followed by uncoating, DNA replication, expression of intermediate and late genes, and culminating in virion maturation and dissemination (36, 37). To pinpoint the step in VACV replication targeted by ZW-2038, we utilized a panel of luciferase reporter viruses in which *Gaussia* luciferase is driven by early (vEGluc), intermediate (vIGluc), or late (vLGluc) VACV promoters. Treatment with ZW-2038 did not affect luciferase activity from the early promoter (**Fig. 3A**), indicating that early gene expression and the replication steps prior to early gene expression are not affected.

**Fig 3.**
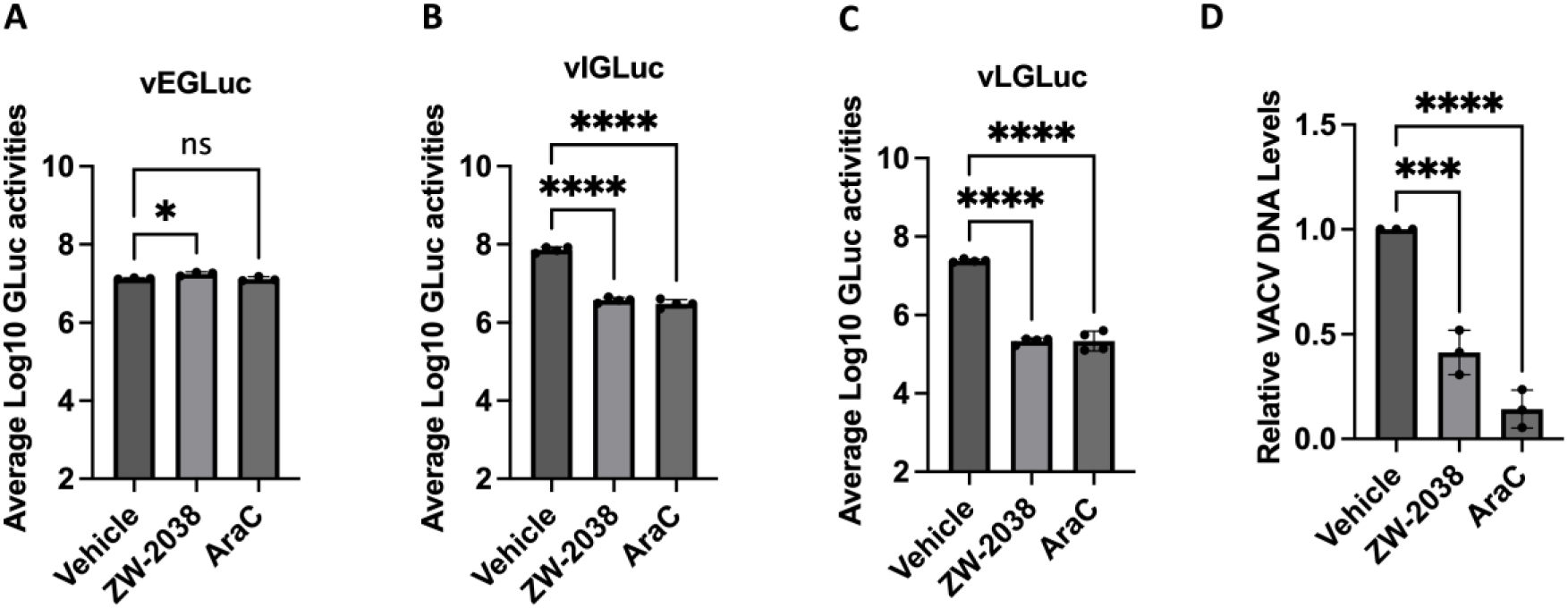
ZW-2038 inhibits VACV gene expression and DNA replication. **(A-C)** HFFs were infected with recombinant VACV expressing *Gaussia* luciferase under the control of early (vEGluc), intermediate (vIGluc), or late (vLGluc) promoters at MOI of 2. Cells were treated with vehicle, 10 µM ZW-2038, or 40 µg/mL AraC. Luciferase activities were measured in the supernatant at 4 hpi for vEGluc and 8 hpi for vIGluc and vLGluc. **(D)** HFFs were infected with VACV at MOI of 2 and treated with either vehicle, 10 µM ZW-2038, or 40 µg/mL AraC. After 8 hours, viral DNA levels were quantified by qPCR using VACV-specific primers. AraC served as a positive control for inhibition of DNA replication. Statistical comparisons were made using one-way ANOVA followed by Dunnett’s test. *P ≤ 0.05; ***P ≤ 0.001; ****P ≤ 0.0001; ns, not significant.

However, reporter activity from intermediate and late promoters was significantly diminished in the presence of ZW-2038, mirroring the suppressive effect of AraC (**Fig. 3B, 3C**). To determine whether this suppression was linked to a defect in viral genome replication, viral DNA levels were quantified at 8 hpi. ZW-2038 caused a marked reduction in viral DNA accumulation, consistent with the replication block observed with AraC (**Fig. 3D**). These findings demonstrate that ZW-2038 suppresses VACV replication by interfering with VACV DNA replication, which in turn prevents the expression of downstream intermediate and late genes.

### ZW-2038 significantly inhibits MPXV and CPXV replication in primary HFFs

Next, we examined the ability of ZW-2038 for its efficiency in inhibiting other orthopoxviruses under a single concentration (10 µM). First, we tested its ability to suppress the replication of MPXV-MA001 2022 isolate in primary HFFs. ZW-2038 significantly suppressed MPXV-MA001 replication in HFFs by over 15-fold MOI of 1 (**Fig. 4A**) and by over 3900-fold at MOI of 0.01(**Fig. 4B**). Cowpox virus (CPXV) is a zoonotic orthopoxvirus that naturally circulates in wild rodents and can infect a broad range of mammals, including humans, causing localized skin lesions and systemic illness, especially in immunocompromised individuals (38). ZW-2038 treatment suppressed the replication of CPXV by 35-fold in HFFs at 48 hpi with MOI of 0.01infection, indicating broad activity of this compound across orthopoxviruses.

**Fig 4.**
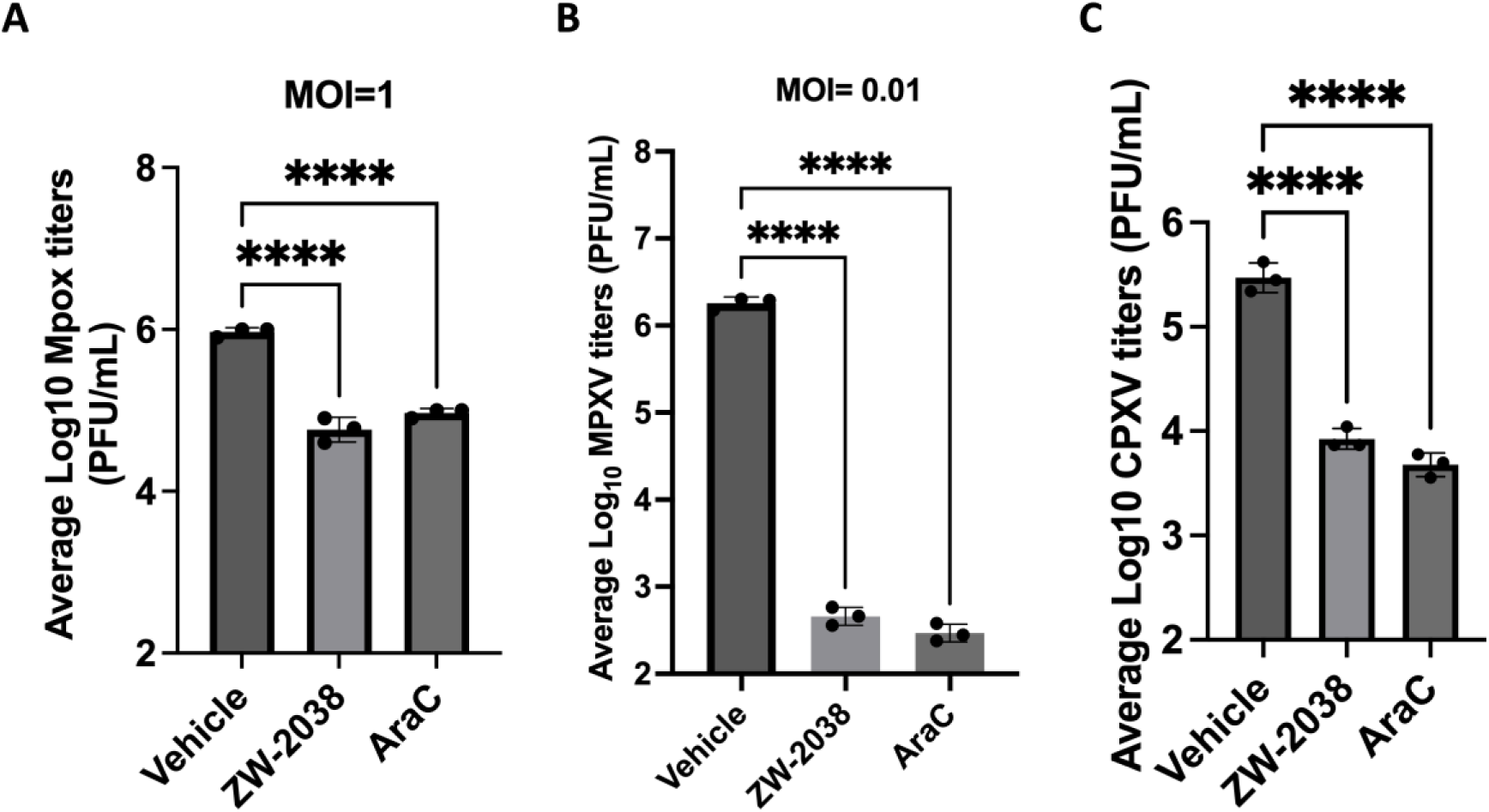
ZW-2038 inhibits MPXV and CPXV replication in primary fibroblasts. **(A)** HFFs were infected with MPXV-MA001 2022 isolate at MOI = 1 and treated with vehicle, 10 μM ZW-2038, or 40 μg/mL AraC at 1 hpi. Virus titers were measured at 24 hpi by standard plaque assay. **(B)** Similar experiments were performed using a lower MPXV MOI = 0.01 to assess multi-cycle replication. **(C)** HFFs were infected with CPXV at MOI = 0.01 and treated with ZW-2038 at10 μM. Viral titers were determined at 24 hpi. In all experiments, ZW-2038 significantly reduced viral replication compared to vehicle control. AraC served as a positive control for replication inhibition. Data represent mean ± SD of biological replicates. Statistical significance was assessed using one-way ANOVA followed by Dunnett’s test. ****P < 0.0001.

### ZW-2038 reduces VACV replication in human and non-human primate enteroids and in *ex vivo* lung tissue models

To assess the antiviral efficacy of ZW-2038 in more physiologically relevant complex systems, we employed intestinal stem cell derived human jejunum and rhesus colon enteroids, which better recapitulate the structural complexity and cellular diversity of the gastrointestinal tract compared to conventional cell lines (39, 40). The enteroids were infected with recombinant VACV expressing *Gaussia* luciferase under a late promoter (vLGluc), and luciferase activity was measured at 24- and 48-hpi. In human jejunum enteroids, vehicle-treated controls showed strong time-dependent increases in luciferase activity, while ZW-2038 treatment significantly suppressed viral replication to a level comparable with AraC (**Fig. 5A**). Similarly, in rhesus colon enteroids, both ZW-2038 and AraC markedly reduced late viral gene expression at 48 hours (**Fig. 5B**).

**Fig 5.**
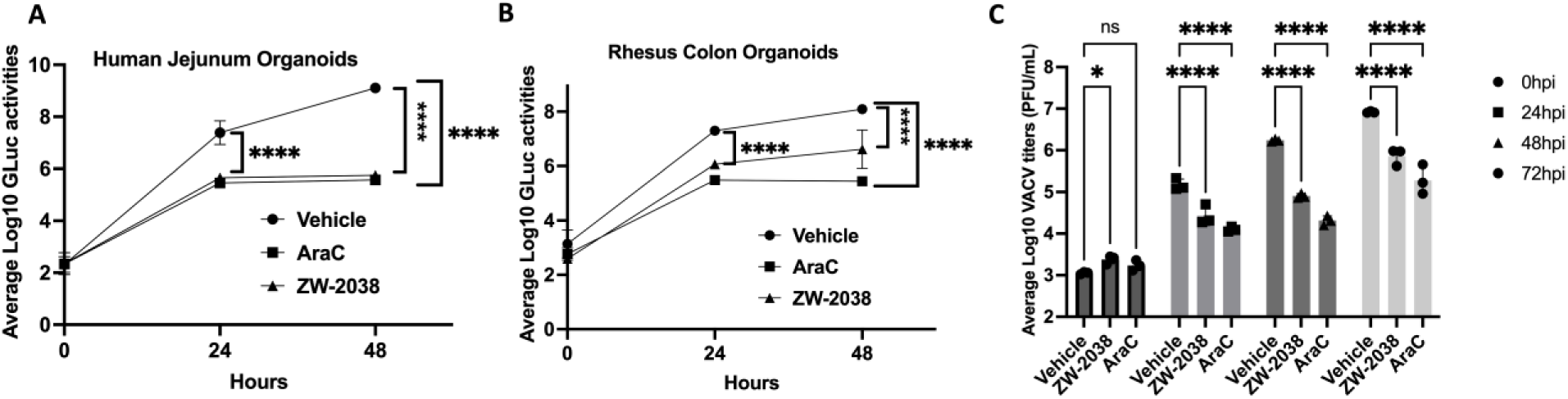
ZW-2038 limits VACV replication in primary enteroids and *ex vivo* lung tissues. **(A–B)** Human jejunum **(A)** and rhesus macaque colon **(B)** enteroids were infected with vLGluc at MOI of 0.01. At 1hpi the media was replaced and treated with either vehicle, ZW-2038 (10 μM), or AraC (40 μg/mL). Luciferase activity was measured from the supernatants at the time of media change (0 hpi), 24, and 48 hpi. ZW-2038 significantly suppressed viral replication in both models. **(C)** Mouse lung tissues were infected with VACV *ex vivo* and treated with ZW-2038 or AraC. Viral titers were quantified at 0, 24, 48, and 72 hpi by plaque assay. ZW-2038 markedly reduced VACV titers at all post-infection time points, similar to AraC. This *ex vivo* lung tissue model was previously used in our lab to assess VACV replication kinetics. Statistical analysis was performed using one-way ANOVA followed by Dunnett’s post hoc test. *P < 0.05; ****P < 0.0001; ns, not significant.

To further confirm these findings in a more intact tissue environment, we used an *ex vivo* mouse lung tissue infection model, which we have previously established and validated for studying poxvirus replication (29). Infected lung tissues were treated with vehicle, ZW-2038, or AraC, and VACV titers were measured by plaque assay at 0, 24, 48, and 72 hpi. ZW-2038 consistently reduced viral titers across all time points, showing similar efficacy to AraC (**Fig. 5C**). These results extend our previous findings and demonstrate that ZW-2038 effectively inhibits VACV replication in both human- and animal-derived physiomimetic models. Together, these data highlight the robust antiviral activity of ZW-2038 under more physiologically relevant conditions and support its potential for further development as a poxvirus therapeutic.

### ZW-2038 does not inhibit MPXV resolvase *in vitro*

ZW-2038 chemically contains a chelating triad capable of binding two divalent metals (**Fig. 6A**), a key pharmacophore feature (**Fig. 6C**) shared by numerous FDA-approved antiviral drugs targeting the RNase H-like (RNHL) viral enzymes (41), all approved HIV-1 integrase strand transfer inhibitors (42), and baloxavir, the FDA-approved influenza PA endonuclease inhibitor (43). Interestingly, MPXV resolvase (Mpr) belongs to the RNHL family and is pharmacologically inhibited by a few metal-chelating chemotypes, as exemplified by dihydroxynaphthyridinone (DHN) analog **1** (25, 44), which bears a similar chelating triad (**Fig. 6B**). Given this similarity, we speculated that Mpr may be the antiviral target of ZW-2038. However, when tested against purified Mpr using our recently developed biochemical assay (25), ZW-2038 showed no Mpr inhibition even at 50 µM, whereas the same concentration of the control compound **1** produced nearly complete inhibition (**Fig. 6D**).

**Fig 6.**
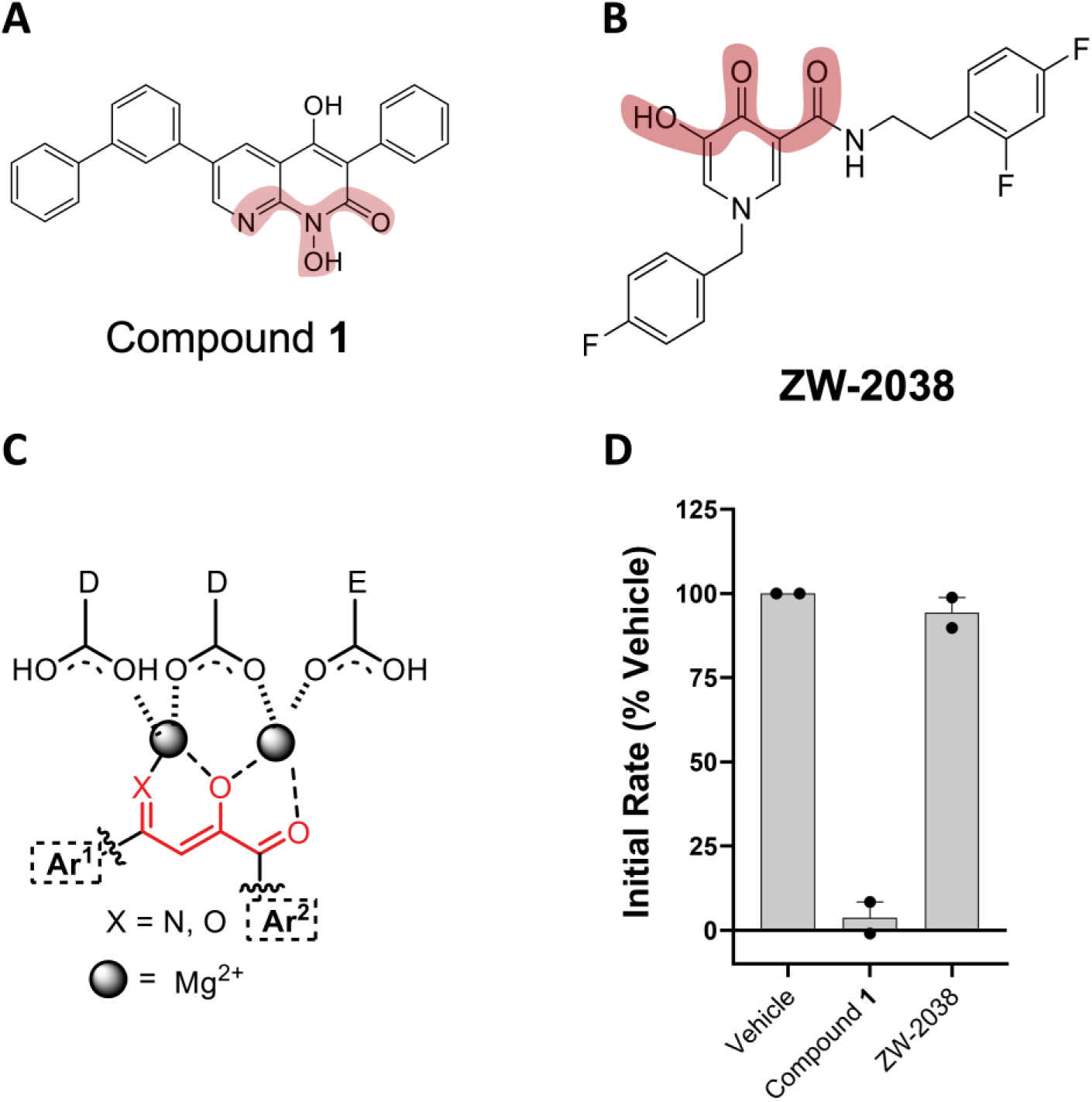
ZW-2038 does not inhibit Mpr *in vitro*. Compound **1** (A) and ZW-2038 (B) both chemically feature a chelating triad (shaded), conforming to a common pharmacophore (C) for inhibiting two-metal dependent viral nucleases. (D) Activity of purified Mpr in the presence of 50 µM Compound **1**, 50 µM **ZW-2038**, or Vehicle (DMSO). Mpr cleavage of a DNA substrate was monitored for 15 min at 37 °C and the rate of product formation was expressed as a percentage of the rate observed with Vehicle treatment. Data are shown as mean ± SEM from two independent experiments performed in duplicate.

### ZW-2038 exhibits low oral bioavailability in mice

To assess the *in vivo* pharmacokinetics of ZW-2038, the compound was administered to CD-1 mice by bolus intravenous (IV) injection and oral gavage. The mean plasma concentration-time profile is presented in **Fig. 7**, and the principal pharmacokinetic parameters are summarized in **Table 1**. Following IV administration, ZW-2038 exhibited a terminal half-life (*t*_1/2_) of 5 h and a large volume of distribution (59 L/kg), consistent with extended systemic retention. After oral dosing, ZW-2038 reached a rapid *T*_max_ of 0.2 h, in line with its previously reported high permeability (24). However, the mean C_max_ achieved orally was only 38 ng/mL at a 20 mg/kg dose, which is 14-fold lower than the IV value of 528 ng/mL obtained with a 10-fold lower dose (2 mg/kg). The oral bioavailability of ZW-2038 was calculated to be 4%, likely reflecting its poor aqueous solubility (2.2 µmol/L or 885 ng/mL) as reported earlier (24).

**Fig 7.**
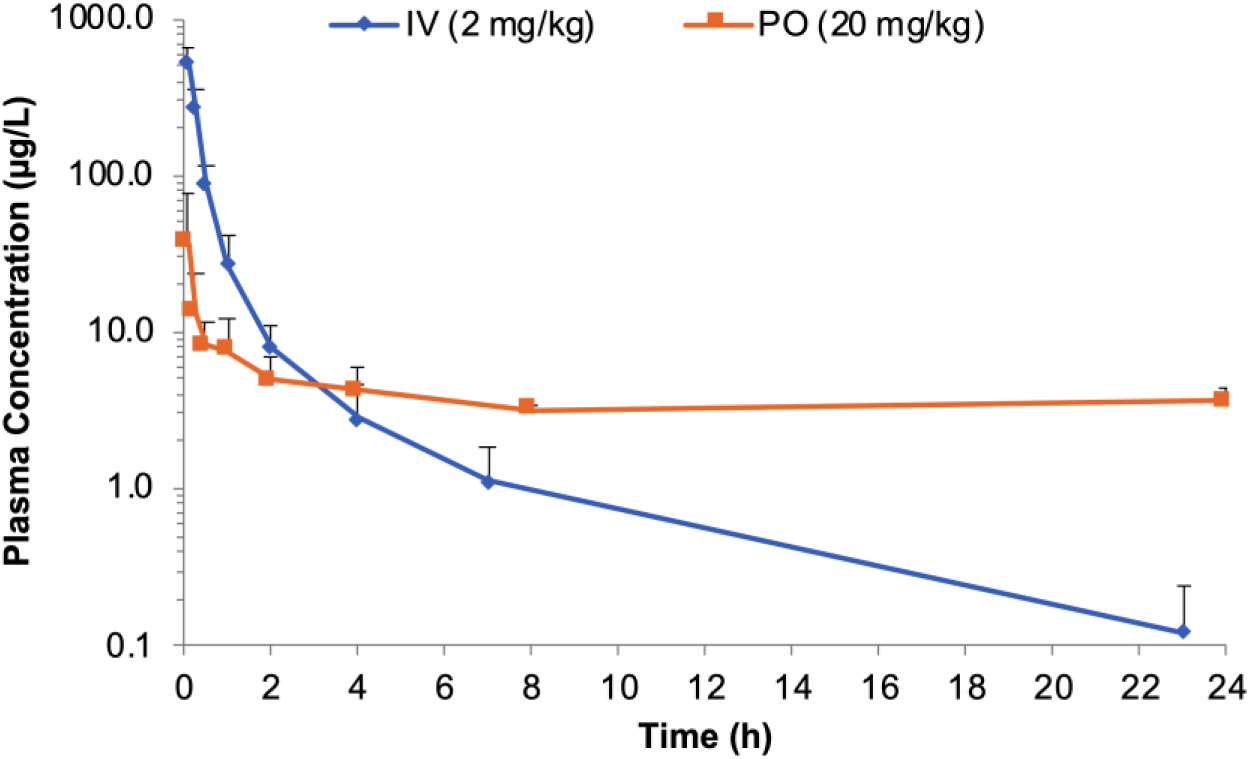
ZW-2038 shows poor oral pharmacokinetics in CD-1 mice. Plasma concentration-time profiles of ZW-2038 following intravenous (IV, 2 mg/kg) and oral (PO, 20 mg/kg) administration in CD-1 mice.

**Table 1.** Key pharmacokinetic parameters of ZW-2038 following intravenous (IV) and oral (PO) administration in CD-1 mice.

| Parameters | IV | PO |
| --- | --- | --- |
| | (2 mg/kg, $n = 4$) | (20 mg/kg, $n = 4$) |
| $t_{1/2}$ (h) | 5 ± 2 | 104 ± 159 |
| $T_{\max}$ (h) | 0.08 ± 0 | 0.2 ± 0.2 |
| $C_{\max}$ (μg/L) | 528 ± 143 | 38 ± 40 |
| AUC<sub>0-t</sub> (h · μg/L) | 238 ± 54 | 98 ± 15 |
| MRT<sub>0-t</sub> (h) | 0.9 ± 0.4 | 11 ± 2 |
| V (L/kg) | 59 ± 33 | 117 ± 46 |
| CL (L/h/kg) | $9 \pm 2$ | $4 \pm 3$ |
| Bioavailability (F, %) | - | 4.1 |

$t_{1/2}$ , terminal half-life. $T_{\max}$ , time to reach $C_{\max}$ . $C_{\max}$ , maximum plasma concentration. $AUC_{0-t}$ , area under the concentration-time curve from the time of dosing to the last quantifiable time point. MRT, mean residence time. V, volume of distribution. CL, clearance.

### ZW-2038 demonstrates negligible hERG channel protein binding *in vitro*

High throughput screening provides a rapid, robust, and cost**-**effective approach for evaluating compound interactions with the hERG channel during early-stage screening. In this study, the Predictor hERG Fluorescence Polarization Assay was used to support lead optimization. The dose response curves for ZW-2038 and the positive control E-4031 are shown in Figure **8**. ZW-2038 exhibited an IC**_50_**of 68 µM, whereas E-4031 demonstrated potent inhibition with an IC**_50_**of 0.0485 µM, consistent with its reported activity range from the manufacturer (0.0095–0.0950 µM). The IC**_50_** of ZW-2038 is approximately 140-fold higher than that of the positive control, indicating very weak *in vitro* affinity for the hERG channel protein.

**Fig. 8.**
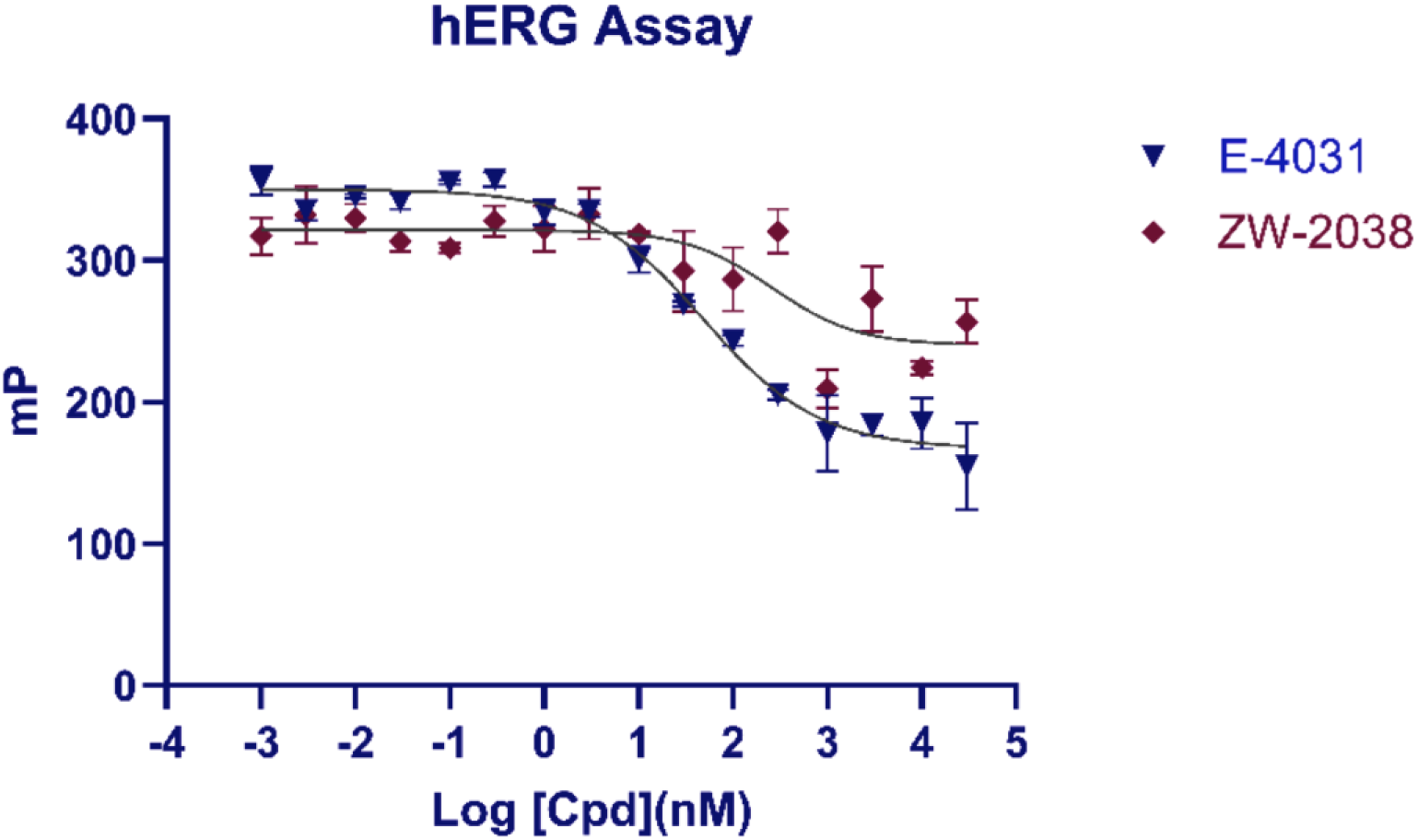
Concentration response curves of ZW-2038 and E-4031 in the hERG assay. The Predictor hERG Fluorescence Polarization Assay was used to measure polarization values of tracer red in presence of ZW-2038 or E-4031. The concentration and corresponding polarization values for ZW-2038(diamonds) and E-4031(circles) were plotted using GraphPad Prism. Non-linear regression was used to calculate IC_50_ values for ZW-2038 (IC_50_ = 68 µM) and E-4031 (IC_50_ = 0.0485 µM). Data represent the mean ± SD of triplicate measurements.

## Discussion

In this work we identified ZW-2038 as a promising antiviral lead against orthopoxviruses. The compound suppressed VACV replication at sub-micromolar concentrations and did not induce cytotoxicity at concentrations of over 500 µM. Mechanistically, ZW-2038 inhibits VACV replication by suppressing VACV DNA replication and downstream intermediate and late gene expression. ZW-2038 exhibits antiviral activity against other orthopoxviruses, including MPXV and CPXV, indicating its broad-spectrum antipoxviral activity. The antiviral effect of ZW-2038 extends beyond the primary cells to human and monkey enteroids and mouse lung tissue models. The activity observed in physiomimetic systems further supports its relevance for translational development.

Although poxviruses are often associated with skin lesions, they can also spread systemically and infect internal organs. Clinical studies have shown that severe mpox cases often involve widespread viral dissemination, including infection of the gastrointestinal (GI) tract (9, 45). Symptoms such as GI inflammation, mucosal damage, and barrier dysfunction have been linked to direct viral replication in the intestine (45–47). Additionally, rectal transmission of MPXV has been reported (48–50). These findings highlight the GI tract as a relevant target tissue during orthopoxvirus infections. Enteroids are lab-grown three-dimensional cultures that closely mimic the structure and function of the intestinal epithelium (39, 51, 52). We, and others (53, 54), have established enteroids derived from both human and animal tissues to study poxvirus-host interactions in the gut model. Because respiratory exposure is a major route of orthopoxvirus transmission and the lung is a key site of viral replication (55, 56), *ex vivo* lung tissue represents a highly relevant model for studying infection and antiviral efficacy. Our findings that ZW-2038 suppresses poxvirus replication in the enteroid models and the *ex vivo* lung tissue model establishes the translational relevance of both our platform and compound.

Unlike DHN analog **1** which potently inhibits poxvirus resolvases (25), ZW-2038 did not inhibit Mpr activity *in vitro*. In a similar manner, while ZW-2038 potently inhibited HCMV it did not inhibit the presumed target pUL89C (24), another RNHL viral nuclease. Taken together, our current studies suggest that ZW-2038 may target another two-metal dependent viral enzyme (57), such as viral exonuclease encoded by both poxvirus and herpesvirus. It is also possible that ZW-2038 interferes with host cell factors required for efficient viral genome synthesis or transcriptional activation. These findings point to a distinct mode of action for ZW-2038 that warrants further investigation to pinpoint its precise molecular target.

ZW-2038 is predicted by Schrödinger QikProp (58) to have good drug-like properties, with hERG blocking being the only potential concern (**Table 2**). In our previous ADME studies (24), ZW-2038 exhibited excellent plasma stability, good microsomal stability and high permeability. However, it also displayed poor aqueous solubility which predicts low oral bioavailability. Although ZW-2038 consistently demonstrated prolonged systemic retention with a large mean residence time in the mice PK studies intended to further characterize ZW-2038, its oral pharmacokinetic profile was more variable compared to IV administration, presumably due to its poor aqueous solubility and limited bioavailability. This necessitates future solubility-driven analog synthesis for lead optimization.

**Table 2.**
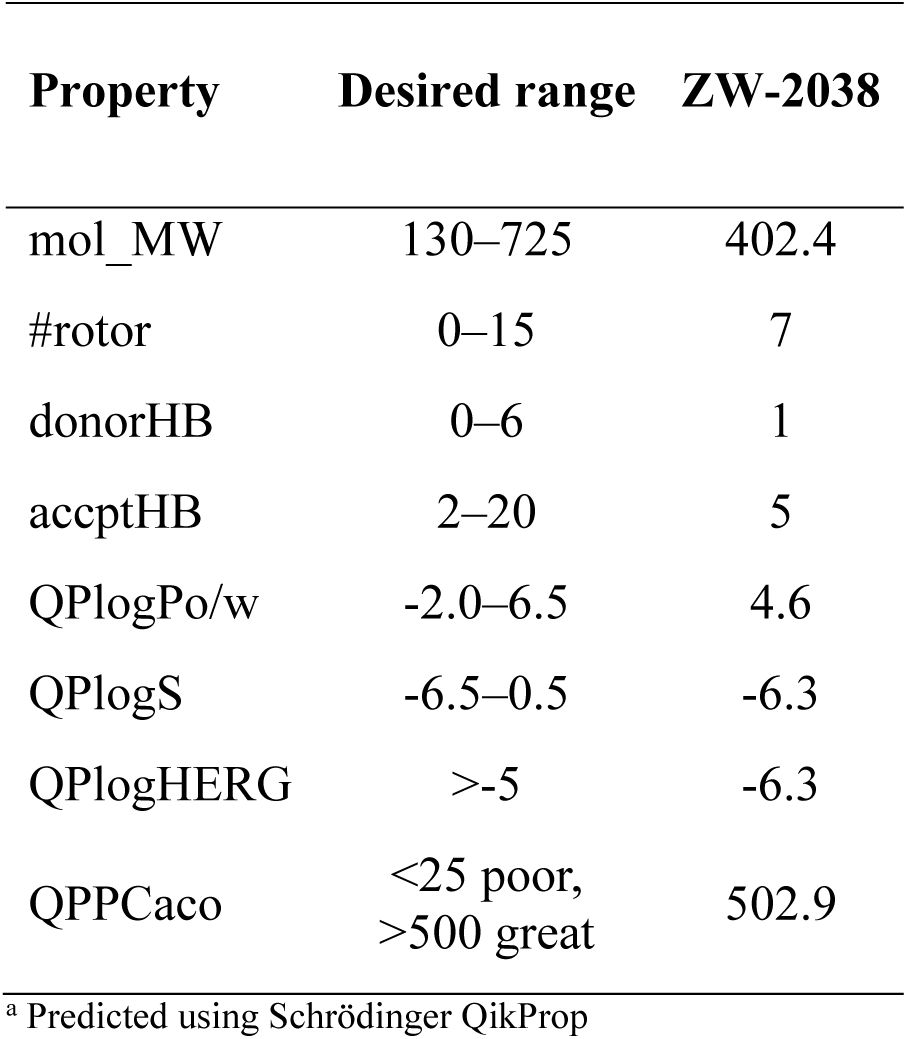
*In silico* physicochemical property profiling of ZW-2038^a^.

During the early stages of drug discovery and development, it is essential to evaluate the safety profile of a new chemical entity and potential adverse effects. Since *in silico* physicochemical property prediction flags hERG as a potential liability for ZW-2038 (**Table 2**), we tested it in our in-house hERG assay. ZW-2038 produced an IC_50_ of 68 µM, which is approximately 140 times higher than positive control E-4031(IC_50_ = 0.0485 µM). This suggests that hERG blocking is unlikely to be a concern for ZW-2038.

In summary, we characterized the antiviral activity of ZW-2038 against VACV, CPXV, and MPXV in primary human fibroblasts and physiologically relevant platforms. We demonstrated promising inhibition of virus replication across multiple orthopoxviruses and infection models, including human and non-human primate enteroids and *ex vivo* lung tissue. We have also identified improving solubility as a main goal for future lead optimization. Structure–activity relationship (SAR) studies will help identify key chemical features driving potency and solubility. The optimized effective compound(s) after SAR optimization will be future evaluated in small animal models of orthopoxvirus infection to determine *in vivo* efficacy, which will pave the way forward to clinical translation for the treatment of mpox and other orthopoxvirus diseases.

## Supporting information

Supplementary information for the synthetic scheme and analytical data

## Acknowledgments

We thank Dr. Nicholas Wallace (Kansas State University) for providing HFFs, Dr. Bernie Moss (National Institute of Health) for various reagents, Dr. Mary Estes (Baylor College of Medicine) for the human enteroid line (HIE J2) and Dr. Hideki Aihara (University of Minnesota) for the Mpr protein.

## Data availability

Data will be made available on request.

