## Supplementary information for the synthetic scheme and analytical data for "Discovery and Characterization of a Hydroxypyridone-3-carboxamide Analog as an Antiviral Lead against Orthopoxviruses"

**Scheme 1.** Synthesis of HPCAm Analog **ZW2038**.

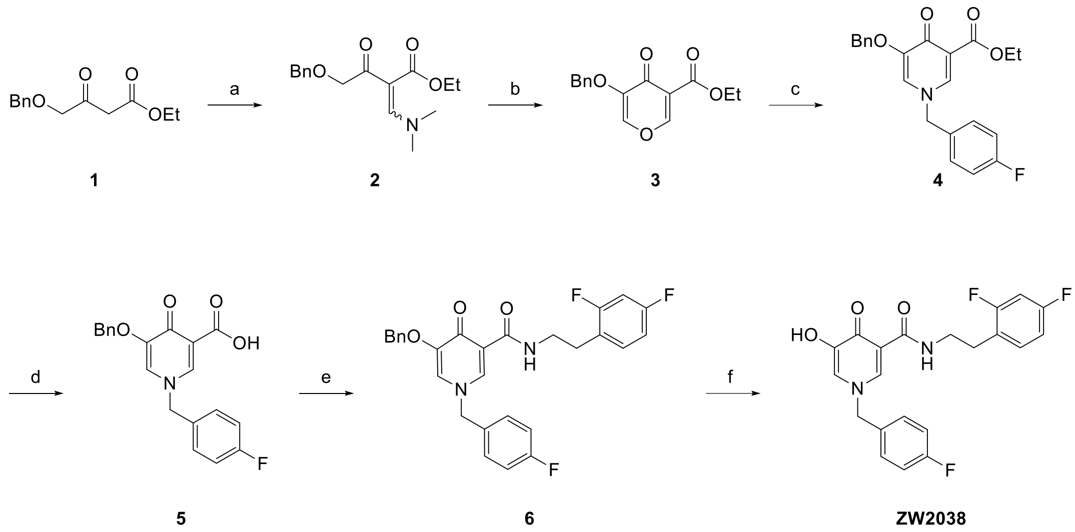


**Reagents and conditions:** a) DMF-DMA, 100 °C, 78%; b) Ethyl formate, KOtBu, THF, rt, 85%; c) 4F-$PhCH_2CH_2NH_2$, EtOH, microwave: 100 °C, 15 min; d) 2M NaOH, EtOH, 90 °C, 4−6h, 68% (two steps); e) 2,4-di $FPhCH_2NH_2$, HATU, DIPEA, DMF, rt, 12h, 71%; f) TFA, microwave: 100 °C, 10 min, 45%.

*N-(2,4-difluorophenethyl)-1-(4-fluorobenzyl)-5-hydroxy-4-oxo-1,4-dihydropyridine-3-carboxamide (**ZW2038**)*

White solid. $^1$H NMR (400 MHz, DMSO-*d*$_6$) $\delta$ 10.18 (t, *J* = 5.8 Hz, 1H), 9.14 (s, 1H), 8.48 (d, *J* = 2.3 Hz, 1H), 7.66 (d, *J* = 2.3 Hz, 1H), 7.49 – 7.43 (m, 2H), 7.37 (td, *J* = 8.6, 6.6 Hz, 1H), 7.28 – 7.21 (m, 2H), 7.18 (ddd, *J* = 10.5, 9.4, 2.6 Hz, 1H), 7.02 (tdd, *J* = 8.5, 2.6, 1.0 Hz, 1H), 5.25 (s, 2H), 3.56 (q, *J* = 6.6 Hz, 2H), 2.84 (t, *J* = 6.9 Hz, 2H). $^{13}$C NMR (101 MHz, DMSO-*d*$_6$) $\delta$ 169.9, 164.6, 162.5 (d, *J* = 245.4 Hz), 161.4 (dd, *J* = 244.9, 12.1 Hz), 161.1 (dd, *J* = 247.4, 12.2 Hz), 149.8, 139.9, 132.9 (d, *J* = 3.0 Hz), 132.5 (dd, *J* = 10.1, 6.5 Hz), 130.8 (d, *J* = 8.1 Hz), 122.8 (dd, *J* = 16.2, 3.0 Hz), 122.6, 116.3 (d, *J* = 22.2 Hz), 115.7, 111.8 (dd, *J* = 21.2, 4.0 Hz), 104.1 (t, *J* = 26.3 Hz), 59.2, 38.9, 28.1. $^{19}$F NMR (400 MHz, DMSO-*d*$_6$) $\delta$ -112.97 (p, *J* = 8.1 Hz), -113.53 (tt, *J* = 9.7, 5.5 Hz), -114.27 (q, *J* = 8.8 Hz). HRMS (ESI+): *m/z* calcd for $C_{21}H_{17}F_3N_2O_3$ $[M+H]^+$ 403.1270; found 403.1253.

[1]H, [13]C, [19]F-NMR, HPLC and HRMS spectra of **ZW2038**

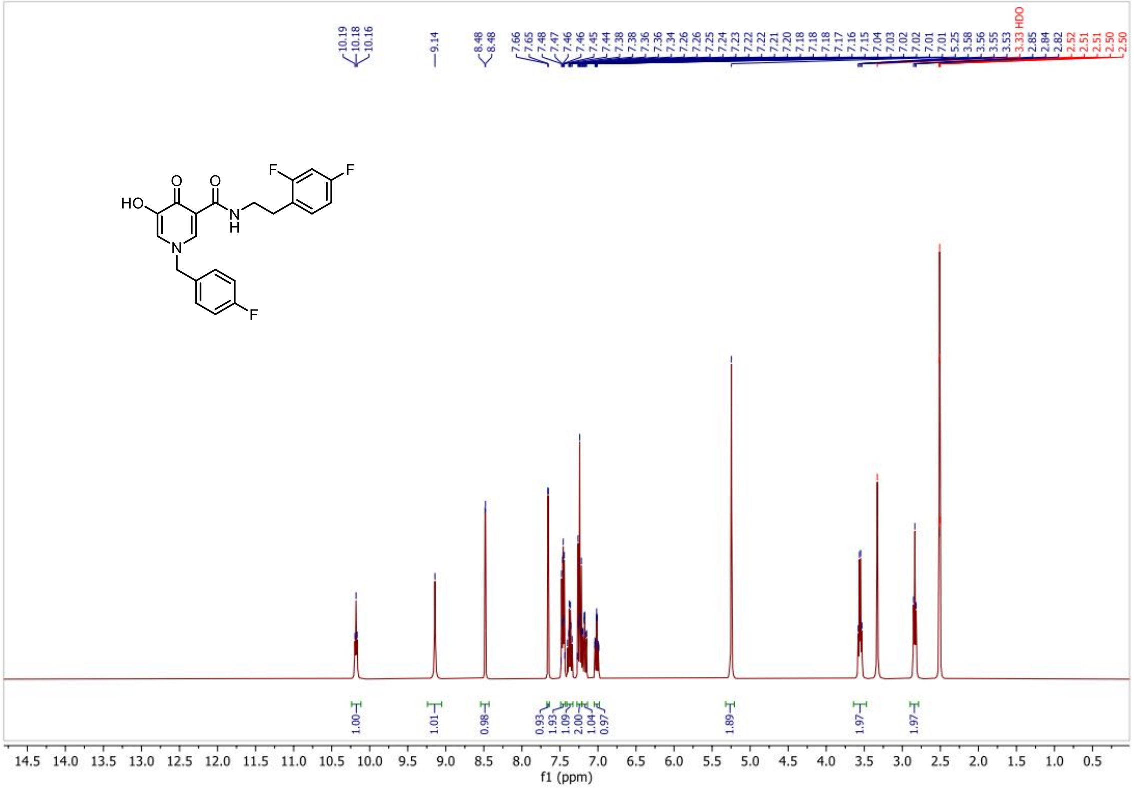


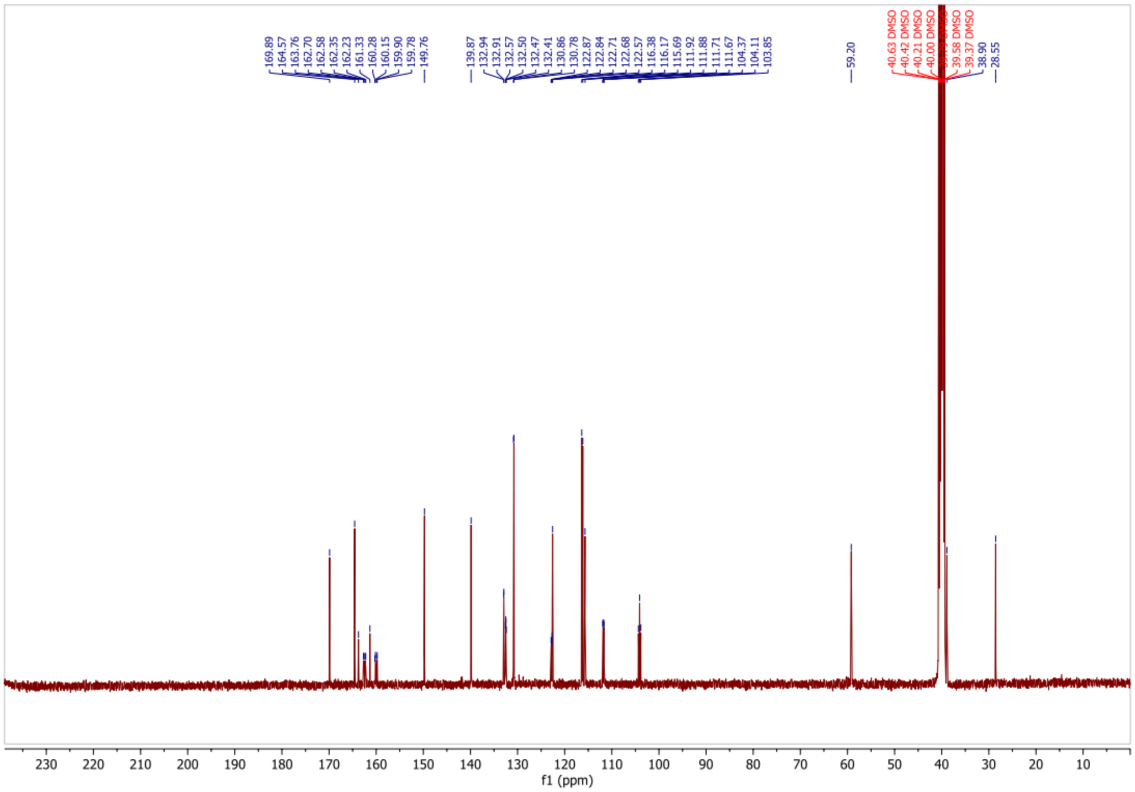

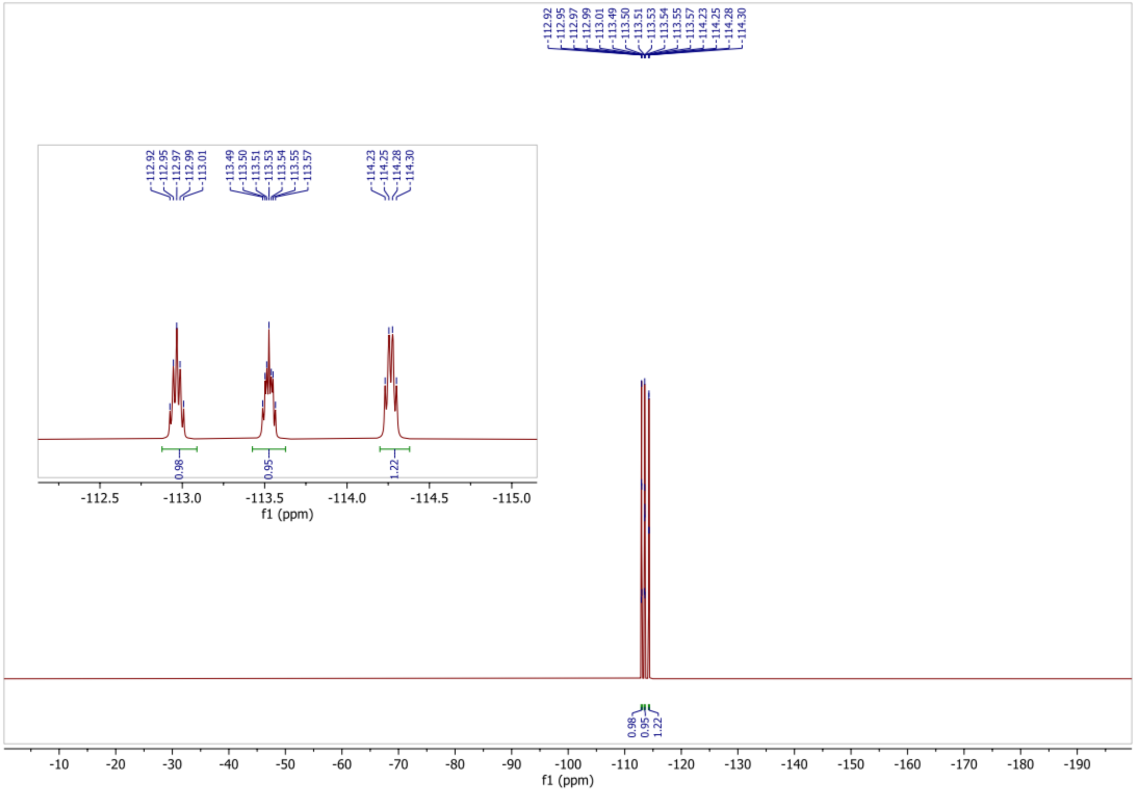


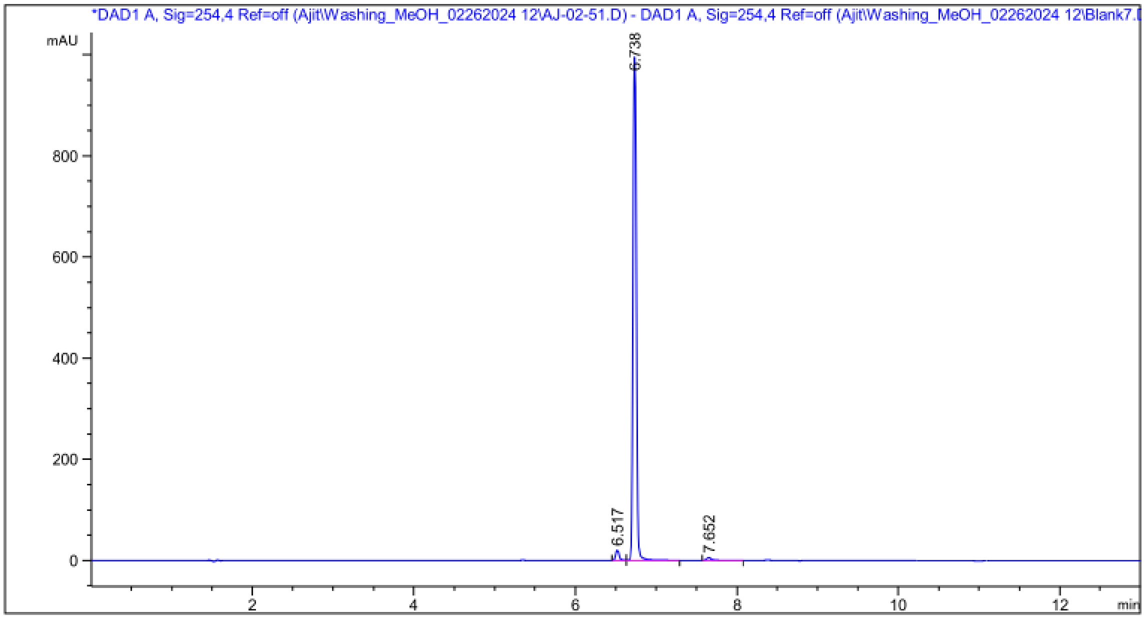


| Peak # | RetTime [min] | Type | Width [min] | Area [mAU*s] | Height [mAU] | Area % |
|---|---|---|---|---|---|---|
| 1 | 6.517 | BV | 0.0489 | 63.95525 | 20.23915 | 2.1619 |
| 2 | 6.738 | VB | 0.0477 | 2872.89941 | 995.06824 | 97.1130 |
| 3 | 7.652 | BB | 0.0608 | 21.45243 | 5.34214 | 0.7252 |

Totals : 2958.30709 1020.64953

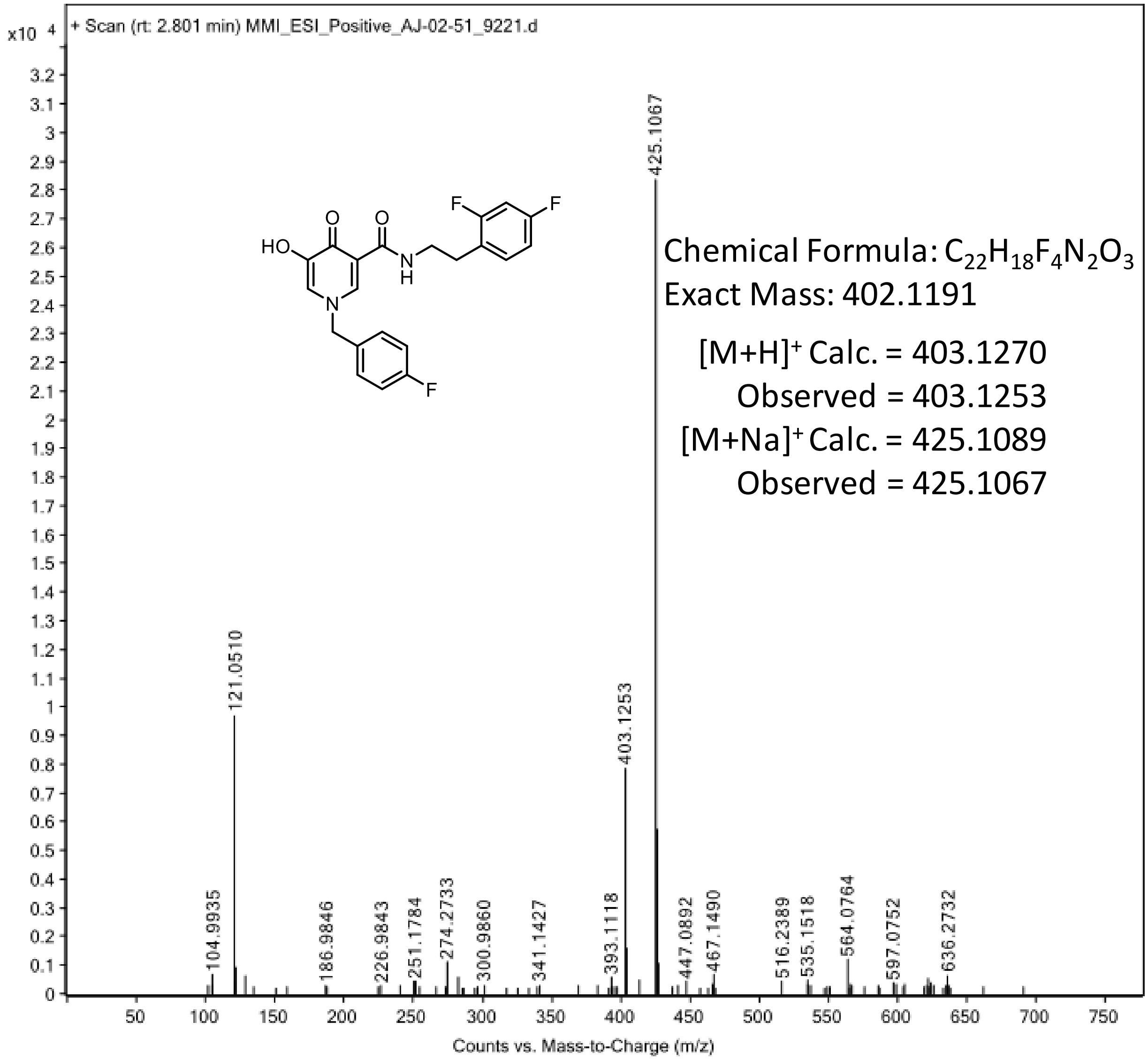

+ Scan (rt: 2.801 min) MMI_ESI_Positive_AJ-02-51_9221.d
x10 4
425.1067
121.0510
403.1253
104.9935
186.9846
226.9843
251.1784
274.2733
300.9860
341.1427
393.1118
447.0892
467.1490
516.2389
535.1518
564.0764
597.0752
636.2732
HO
O
O
N
H
N
F
F
F
Chemical Formula: C22H18F4N2O3
Exact Mass: 402.1191
[M+H]+ Calc. = 403.1270
Observed = 403.1253
[M+Na]+ Calc. = 425.1089
Observed = 425.1067
Counts vs. Mass-to-Charge (m/z)
